# Pick-up Single-Cell Proteomic Analysis for Quantifying up to 3000 Proteins in a Tumor Cell

**DOI:** 10.1101/2022.06.28.498038

**Authors:** Yu Wang, Zhi-Ying Guan, Shao-Wen Shi, Yi-Rong Jiang, Qiong Wu, Jie Wu, Jian-Bo Chen, Wei-Xin Ying, Qin-Qin Xu, Qian-Xi Fan, Hui-Feng Wang, Li Zhou, Jian-Zhang Pan, Qun Fang

**Author notes:** These authors contributed equally: Yu Wang, Zhi-Ying Guan. **Correspondence and requests for materials** should be addressed to Q.F.

## Abstract

The shotgun proteomic analysis is currently the most promising single-cell protein sequencing technology, however its identification level of ∼1000 proteins per cell is still insufficient for practical applications. Here, we develop a pick-up single-cell proteomic analysis (PiSPA) workflow to achieve a deep identification capable of quantifying up to 3000 protein groups in a tumor cell using the label-free quantitative method. The PiSPA workflow is specially established for single-cell samples mainly based on a nanoliter-scale microfluidic liquid handling robot, capable of achieving single-cell capture, pretreatment and injection under the pick-up operation strategy. Using this customized workflow with remarkable improvement in protein identification, 1804-3349, 1778-3049 and 1074-2487 protein groups are quantified in single A549 cells (*n* = 37), HeLa cells (*n* = 44) and U2OS cells (*n* = 27), respectively. Benefiting from the flexible cell picking-up ability, we study tumor cell migration at the single cell proteome level, demonstrating the potential in practical biological research from single-cell insight.

---

Nowadays, single-cell genomic^1^ and transcriptomic technologies^2^ have been well developed. However, the development of the single-cell proteomic technology faces major technical challenges, because the protein content in a single cell is extremely low and proteins are difficult to be amplified as nucleic acids. So far, a variety of protein analysis techniques at the single-cell level have been developed, including those based on specific antibody labeling such as flow cytometry^3^, fluorescence imaging^4^, Western blotting^5^ and mass cytometry^6^, as well as those based on non-labelling proteome analysis technique with high-resolution mass spectrometry (MS). At present, among the proteomic analysis techniques based on MS, the shotgun technique using the bottom-up strategy usually demonstrates the maximum depth and breadth of protein identification^7^, with which usually 5000-7000 proteins^8-10^ can be identified in a single measurement with a liquid chromatography-mass spectrometry (LC-MS) system for a sample containing a large number of cells with a protein amount in the range of 200-1000 ng. Therefore, in recent years, a variety of single-cell proteome analysis approaches based on the shotgun technique have been developed^11,12^.

A typical single-cell shotgun proteomic analysis process includes the sorting of target cells from a large number of sample cells, the pretreatment of single-cell samples, the LC injection and separation, and ESI-MS/MS detection of the digested peptides. The sample pretreatment process includes multi-steps of cell lysis, protein reduction, alkylation, enzymatic digestion and termination of the digestion. For such single-cell samples with extremely small protein amounts (ca. 100-500 pg per cell^13^), if the above-mentioned series of sample pretreatment operations are carried out with a microliter-scale reactor such as a centrifuge tube^11^ or a multi-well plate^14^, obvious sample loss will occur during the sample pretreatment and transferring process, which will significantly reduce the number of protein identification and thus severely limit the identification depth of single-cell proteomics.

To break through the barriers of the identification depth of single-cell proteomics, one strategy to address the above challenge is to perform the sample pretreatment in nanoliter-scale in-situ microreactors, such as the oil-air-droplet (OAD) chip^15^, the integrated proteome analysis device (iPAD)^16^ and the nanodroplet sample preparation (nanoPOTS) platform^12,17-20^. Compared with the conventional microliter-scale reactors used in routine laboratories, these microreactors have a volume reduction of hundreds of times, which can avoid the excessive dilution of the trace amounts of single-cell samples, effectively improve the reaction efficiency and reduce the sample loss caused by the adsorption of the sample components on the reactor surface during the pretreatment process. Based on these methods, up to 300-1100 proteins were able to be identified from single cells with label-free approach^12,16,17,19^. With the tandem mass tag (TMT) approach, the maximum up to 1500 proteins were quantified from single acute myeloid leukemia (AML) cells^21^. However, most of the reported nanoliter-scale microreactors required to use microfabricated microchips or devices as the microreactors, and needed special sample injection devices and additional operations to complete the injections of the nanoliter-volume samples. Currently, an important emerging trend is to develop single-cell proteomic platforms capable of integrating whole-process operations for promoting the practicality and popularization of single-cell proteomic analysis. Some integrated platforms were developed by using the combination of commercial instruments and self-developed systems (such as the autoPOTS platform^22^ and the T-SCP platform^23^) or an integrated microfluidic chip^24,25^, to complete the whole process of single-cell proteomic analysis, with 300-1500 proteins identified from single cells.

In spite of the significant progresses obtained in single-cell proteomics, how to further improve the protein identification depth and simultaneously simplify the device and operation to achieve practical whole-process proteomic analysis at the single-cell level still presents great challenges. Here, we developed a total workflow solution for single-cell proteomic analysis (Fig. 1) capable of achieving deep identification of more than 3000 protein groups in a single tumor cell. More automatic and convenient operation was performed by using a probe-based microfluidic liquid handling robot coupled with a commercial liquid chromatograph (LC) and a trapped ion mobility spectrometry (TIMS) QTOF mass spectrometer. The automated pick-up operation mode based on capillary probes was adopted throughout the pick-up single-cell proteomic analysis (PiSPA) workflow, including the sorting of single cells and multi-step single-cell pretreatment to digest cellular proteins into peptides in nanoliter reactors, as well as the injection of the peptide samples to the LC column. In addition, we utilized a single-cell customized strategy that fully considered the effects of the unique properties of single cells *vs*. bulk cells on sample pretreatment, separation, and detection to establish the series of measures throughout the PiSPA workflow. This strategy enables a much deeper depth of protein identification in single-cell analysis than those previously reported in the literatures. We applied this platform in the single-cell proteomic analysis of three kinds of tumor cells, HeLa, A549 and U2OS cells, as well as the single-cell proteomic study of migrating tumor cells.

**Fig. 1.**
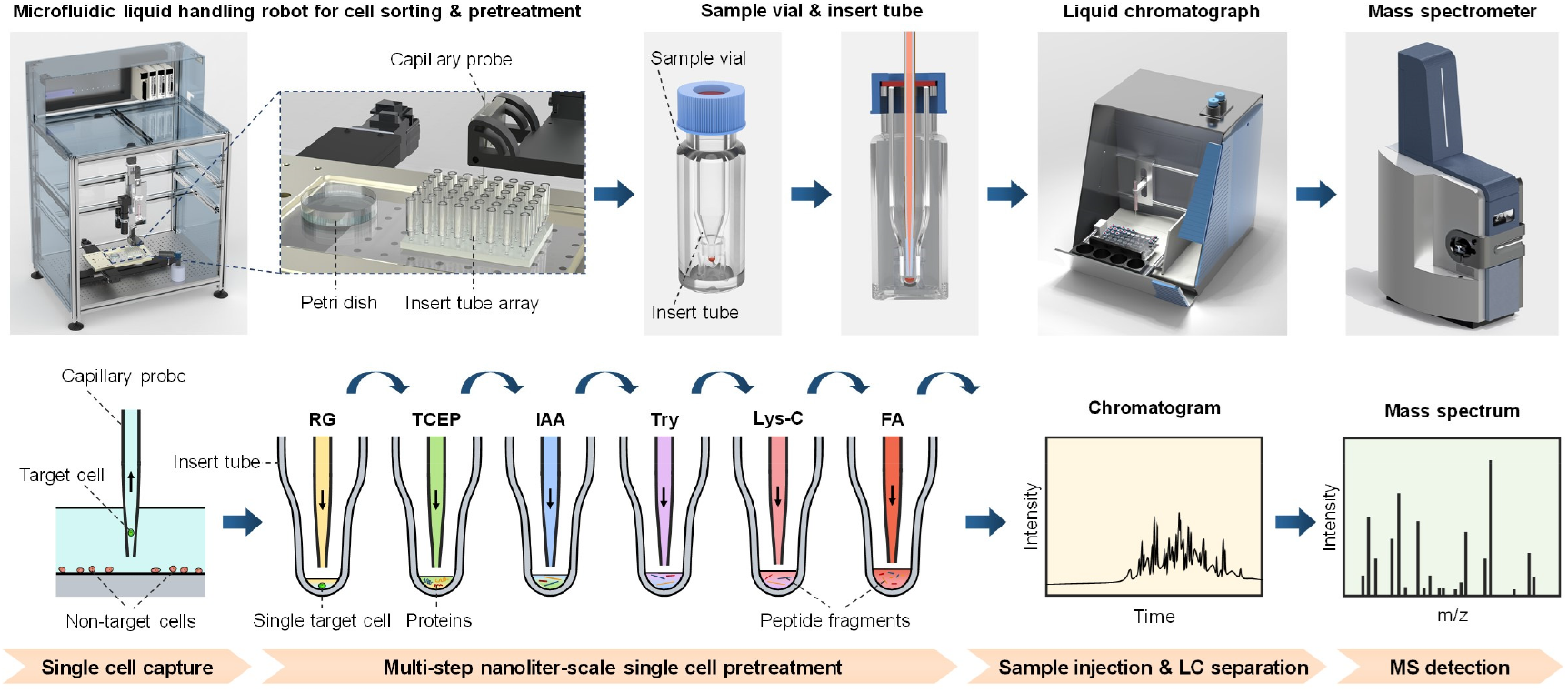
Schematic diagram of the PiSPA workflow for single cell proteomic analysis. The PiSPA workflow was conducted using a probe-based microfluidic liquid handling robot for cell sorting and pretreatment, a commercial LC system with an autosampler, and a tims-QTOF mass spectrometer. The microfluidic liquid handling robot completed the sorting of single cells and the multi-step pretreatment of the single cell samples with the automated pick-up operation mode, including nanoliter-scale cell lysis, protein reduction, alkylation, enzymatic digestion and termination of the digestion. Insert tubes coupled with sample vials were used as the nanoliter microreactors for sample pretreatment of single cells. After sample pretreatment, the insert tubes & sample vials were used as sample tubes for the autosampler of the LC system to perform the sample injection, LC separation and subsequent MS detection of the digested peptide components from single cells.

## Results

### Establishment of the PiSPA workflow

We first used an improved microfluidic liquid handling platform^26^ in the picoliter to nanoliter range previously developed by the authors’ group based on the sequential operation droplet array (SODA) technique^27^. It achieved the automated bright-field or fluorescence imaging and capture of single target cells from cell suspensions by a tapered capillary probe connected with a high-precision syringe pump under the probe pick-up mode. Under the optimized conditions, multiple types of cells such as HeLa, A549, U2OS cells could be captured with a capture success rate of 95%.

After the probe captured the target cell, we controlled the platform to dispense the cell into a commercial insert tube instead of using a microchip as the microreactor as previously reported^15^. The insert tubes can be seamless compatible with the autosampler of commercial LC-MS systems, without the need for microfabricated microchips and troublesome sample injection operations^14,15,19^. However, even for the smallest insert tubes available on the market, their volumes are still hundreds of microliters, which are far from a nanoliter-scale microreactor required for single-cell sample pretreatment. We solved this difficulty using the conical bottom tip of the insert tubes to load the nanoliter reaction solution. In addition, the evident evaporation effect of the nanoliter reaction solutions during the heating reaction process was suppressed by adding 100 μL of water and a sealing cap to the sample vial outside the insert tube to form an internal high-humidity environment. With the precise manipulation ability for nanoliter-scale liquids, the single-cell capture platform was continuously adopted to complete the multi-step nanoliter-scale reagent addition operation to sequentially achieve cell lysis, protein reduction, alkylation, digestion and reaction termination, under the pick-up operation mode. We performed a comparison experiment to conduct sample pretreatment of single HeLa cells with reaction volumes of microliters (5 μL) and nanoliters (∼400 nL). The protein identification number of the nanoliter reactors was double that of the microliter reactors (Supplementary Fig. 1). The advantage of nanoliter-scale reactors in single cell proteomic analysis had also been demonstrated in many previous literatures^13,16,20^.

Before performing single-cell proteomic analysis of tumor cells, we first tested the repeatability of the LC-MS system under the optimized conditions by running 10 consecutive analysis of 200 pg of standard HeLa cell digestion sample in data independent acquisition (DIA) and data dependent acquisition (DDA) modes. On average, 1469 and 618 protein groups (Fig. 2a) with variation coefficients of 4.7% and 2.3% were stably quantified under the DIA and DDA mode, respectively. The pairwise analysis of arbitrary two replications showed that all of the Pearson correlation coefficients were over 0.99 (DIA mode) and 0.85 (DDA mode) (Fig. 2b), demonstrating the good stability (especially in DIA mode) of the system in the proteomic analysis at the single-cell level.

**Fig. 2.**
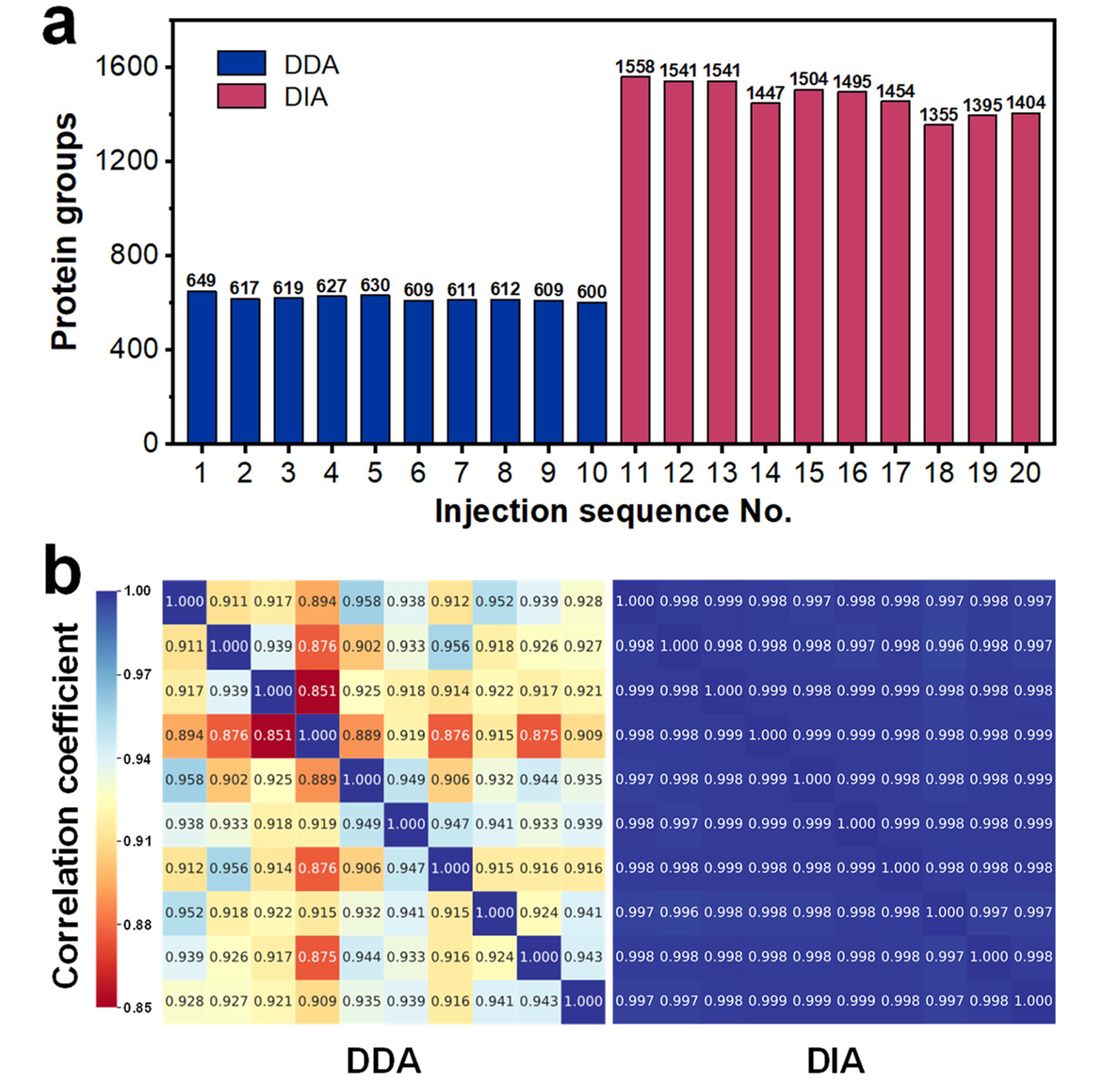
System optimization and performance. Test for the repeatability of 10 consecutive analysis of 200 pg standard HeLa cell digestion sample using the LC-MS system in DIA and DDA modes. The results of protein identification number under the DIA and DDA mode (**a**) and their Pearson correlation coefficients of pairwise analysis (**b**) are shown.

### Single-cell proteomic analysis of tumor cells

Using the present workflow and platform, we performed single-cell proteomic analysis of three types of tumor cells, A549, HeLa, and U2OS cells, using the DIA and DDA modes. Under the DIA mode, an average of 2467 (1804-3349), 2421 (1778-3049) and 1705 (1074-2487) protein groups were quantified in single A549 cells (*n* = 37), HeLa cells (*n* = 44) and U2OS cells (*n* = 27), respectively (Fig. 3a). Using the match between runs (MBR) algorithm, an average of 3008 (2449-3500), 2926 (2278-3257) and 2259 (1621-2904) protein groups were quantified in the same samples of single A549, HeLa and U2OS cells, respectively. Under the DDA mode at the same analysis conditions, an average of 1328 (712-2129), 1290 (664-2198) and 1005 (536-1519) protein groups were quantified in single A549 cells (*n* = 56), HeLa cells (*n* = 68), and U2OS cells (*n* = 24) (Fig. 3a).

**Fig. 3.**
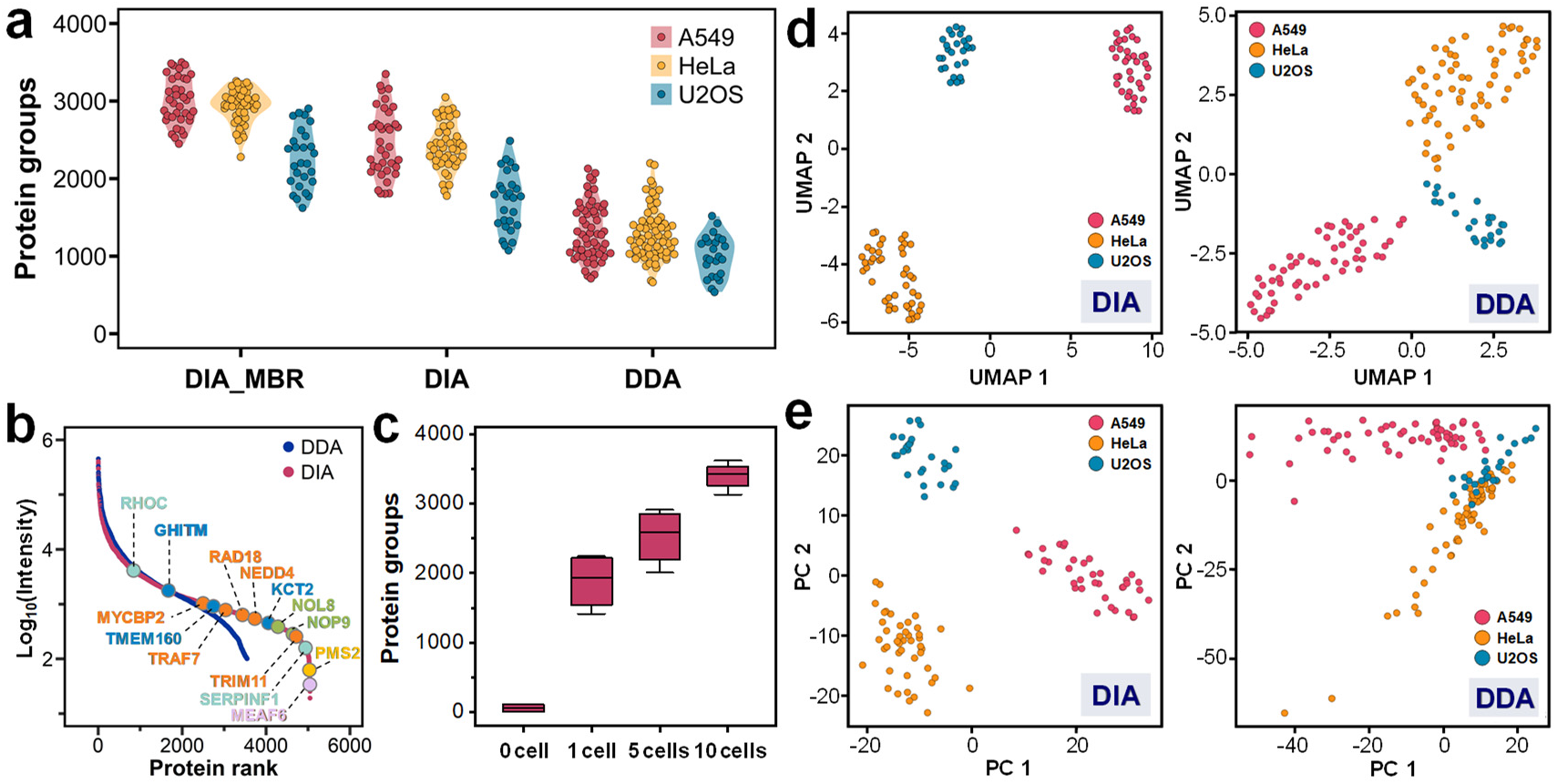
Application of the PiSPA platform in proteomic analysis of single tumor cells. **a**, Quantified protein group numbers of three types of tumor cells (A549, HeLa, and U2OS cells) under the DIA and DDA modes. Under the DIA mode, an average of 3008, 2926 and 2259 protein groups were quantified in single A549 (*n* = 37), HeLa (*n* = 44) and U2OS (*n* = 27) cells with MBR algorithm, and average of 2467, 2421 and 1705 protein groups were quantified in the same single A549, HeLa and U2OS cells without MBR. Under the DDA mode, an average of 1328, 1290 and 1005 protein groups were quantified in single A549 (*n* = 56), HeLa (*n* = 68) and U2OS (*n* = 24) cells. **b**, Rank order of protein abundance in single HeLa cells under the DIA (red dots) and DDA (blue dots) modes. Some of the proteins only quantified in the DIA mode are marked with colored points. **c**, Quantified protein numbers from 0, 1, 5 and 10 HeLa cells with the same procedures and conditions under the DIA mode. **d**,**e** Comparison of UMAP dimensionality reduction analysis (**d**) and principal component analysis (PCA) (**e**) on the measured A549, HeLa, and U2OS cells under the DIA and DDA modes.

Benefiting from the ability of the present platform to flexibly and accurately control the cell number in samples, we tested samples containing 0, 1, 5, and 10 HeLa cells with the same procedures and conditions, and quantified 57, 1880, 2528, and 3399 protein groups (*n* = 4) under the DIA mode, respectively (Fig. 3c). Compared with the single-cell result, further increasing the number of cells to 5 or 10 cells non-linearly increased the protein identification number by about 35% to 80%. These results showed that the present protocol is a customized one that is more suitable for single-cell samples to achieve high identification depths.

We implemented a uniform manifold approximation and projection (UMAP) dimensionality reduction on the above data, three types of single-cell samples could be automatically and clearly clustered (Fig. 3d), reaching recognition accuracies of 100%.

Under the DDA mode, the label-free single-cell protein identification numbers obtained in this work (Fig. 3a) are higher than most of the literature reported results for the same type of cells (e.g. HeLa cells)^12,15-19,22^, which shows the advantage of the present workflow and platform in the identification depth of single-cell proteome analysis. Compared with the single-cell proteomic data set obtained under the DIA mode, the single-cell protein identification numbers based on the DDA mode are relatively low. Under the DIA mode, a total of 5093, 5048 and 4286 protein groups were cumulatively quantified in single A549 (*n* = 37), HeLa (*n* = 44) and U2OS (*n* = 27) cells, while only 3286, 3219 and 2357 protein groups were cumulatively quantified in single A549 (*n* = 56), HeLa (*n* = 68) and U2OS (*n* = 24) cells under the DDA mode (Supplementary Fig. 2). Under the DIA and DDA modes, the ranges of protein abundance spanned nearly 5 and 4 orders of magnitude, respectively. Some important but low abundant proteins could only be quantified under the DIA mode (Fig. 3b), such as mismatch repair endonuclease PMS2^28^ and E3 ubiquitin-protein ligase NEDD4^29^. In addition, although the DDA-based data set could also be distinguished by UMAP, it could not be clearly distinguished into three types of cells with principal components analysis (PCA) as the DIA based data set (Fig. 3e). This may be because the DIA mode enables unbiased acquisition of MS2 information for all the precursors, and the machine learning based DIA-NN algorithm can improve the fragment signal pattern identification and its matching to the precursor level^23^. Both are beneficial to reduce the identification bias of proteins with relatively low abundance in single cell samples, resulting in more comprehensive identification results of single cell proteome.

Cell migration is a common biological process, which has important significance for the study of tumor migration, wound healing, embryonic development, immune response, etc^30^. At present, the scratch assay is frequently used to test the invasion and metastasis ability of adherent tumor cells^31^, while so far there is no report on the study of individual tumor cells with different migration behaviors at the deep-coverage proteome level.

We used HeLa cells to perform the scratch assay, and 9 single cells with significant migration behaviors and 10 single cells without obvious migration behaviors were captured and analyzed with the PiSPA platform.

The data sets of the migrating and non-migrating HeLa cells were analyzed using the partial least squares discriminant analysis (PLS-DA) methods (Fig. 4b). The results show that the protein compositions of the migrating and non-migrating cells have significantly differences, and these proteomic differences are further quantitatively displayed in the volcano plots (Fig. 4c) and hierarchical cluster analysis (HCA) plots (Fig. 4a). After the two-side *t*-test, a total of 143 differential proteins were screened out between the two data sets (*p* < 0.05, fold change > 2). Among them, 32 proteins are up-regulated and 111 proteins are down-regulated in the migrating cells compared to the non-migrating cells.

**Fig. 4.**
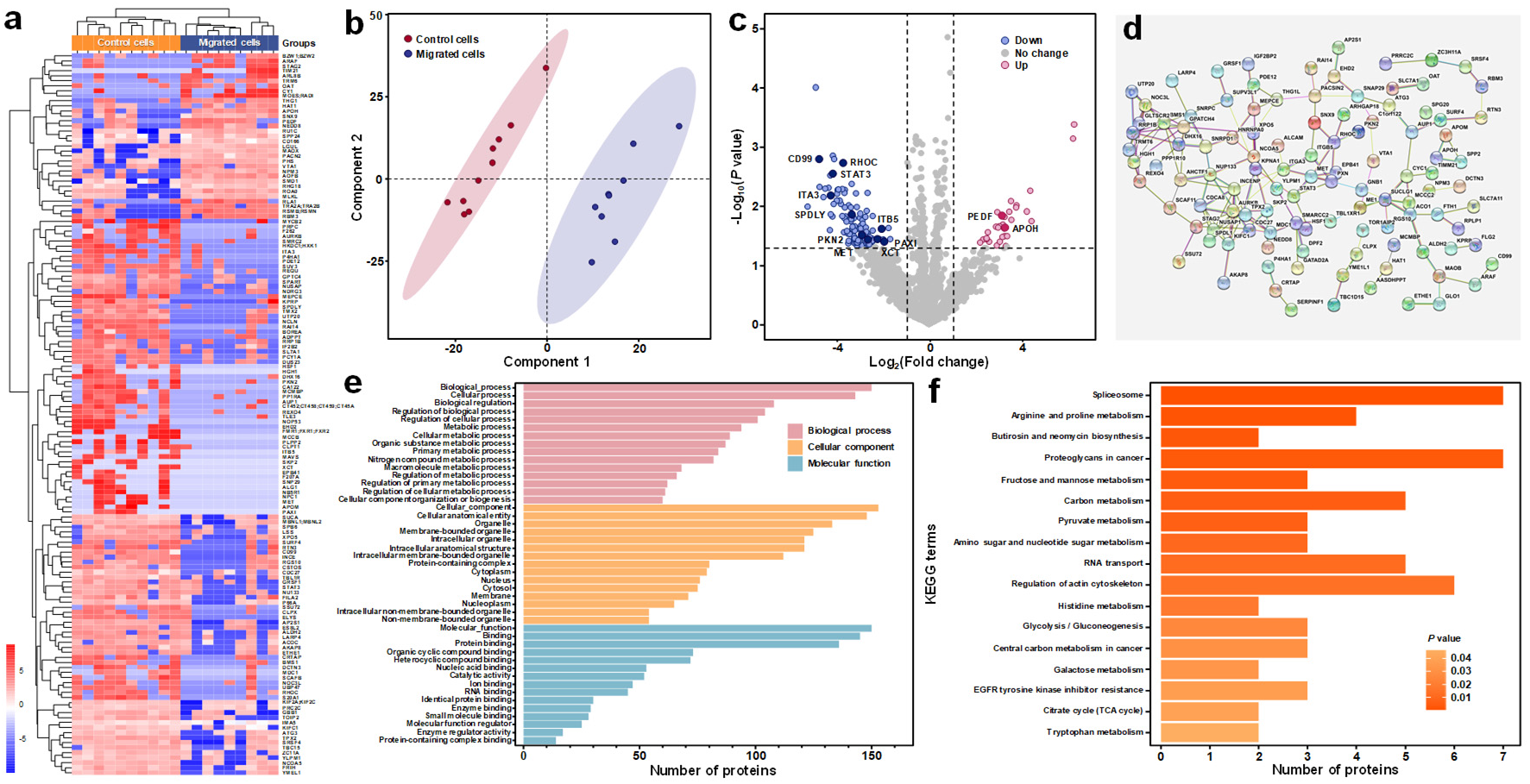
Application of the PiSPA platform for proteomic analysis of single migrated cells in scratch assay. In the scratch assay, 9 single HeLa cells with significant migration behaviors and 10 single HeLa cells without obvious migration behaviors as control cells were captured and analyzed in the experiment of scratch assay. **a**, Hierarchical clustering heatmap of differential proteomes (*p* < 0.05 and fold change > 2) of the tested migrated and control HeLa cells. **b**, Partial least squares discriminant analysis (PLS-DA) of the migrated and control HeLa cells. **c**, Volcano plot showing 32 up-regulated proteins and 111 down-regulated proteins screened out in the migrated cells compared to the control cells, including 12 proteins related to cell migration (marked with dark blue and dark red dots). **d**, Interaction network map of differential proteins according to the String database. **e**,**f**, Enrichment analysis of 143 differential proteins (*p* < 0.05 and fold change > 2) with the Gene Ontology (**e**) and KEGG database (**f**).

Using gene annotation enrichment analysis on Gene Ontology (GO)^32^ terms (Fig. 4e) (*p* < 0.05), 143 differential protein groups were enriched, most of which have important biological functions related to cell migration, location, metabolism, binding, cell cycle, transport and endocytosis. Among them, 12 proteins relate to cell migration, including RHOC^33^, MET^34^, CD99^35^, ITA3, PEDF^36^, STAT3^37^, APOH, ITB5, PAXI, PKN2^38^, SPDLY and XCT^39^. We also enriched 155 signal pathways with the Kyoto Encyclopedia of Genes and Genomes (KEGG)^40^ database, and some of these pathways were reported to be involved in cell migration process, such as the pathways of proteoglycans in cancer^41^, amino sugar and nucleotide sugar metabolism^42^, and glycolysis/gluconeogenesis^43^ (Fig. 4f). Based on the String database^44^, we also obtained the interaction network map of these differential proteins (Fig. 4d).

Although the scratch assay is a frequently-performed biological experiment, the protein variation information at the single cell level during cell migration is largely absent. The PiSPA platform has the ability to observe the behavior of individual migrated cells, reliably pick the target cells up, and achieve deep-coverage proteomic analysis to these cells at the single-cell level to highlight the individual protein differences between cells with different apparent migratory properties. Such a platform has the potential to reveal the intrinsic control factors behind the apparent behaviors that were previously difficult to detect using the conventional omics approaches for large amounts of cell samples, and could provide an effective tool for the cell migration studies as well as the development of anticancer approaches.

The present application of single-cell proteomics in cell migration experiment is just a preliminary proof-of-principle attempt, and similar approaches can be extended to many conventional biomedical experiments, such as susceptibility experiments of anti-tumor drugs.

## Discussion

The PiSPA workflow developed using the nanoliter-scale pick-up operation and the single-cell customized strategy provides a complete set of solution for deep-coverage single-cell proteomics analysis. The platform requires relatively simple equipment and is easy to operate and couple with currently-available commercial instruments.

Considering that the conventional proteomic analysis of samples containing thousands of cells can typically identify about 5000-7000 protein groups^8,24^, our PiSPA platform has the ability to quantify nearly half this level for a single tumor cell with a significant improvement over many current single-cell proteomic analysis systems. This may imply that the single-cell proteomics research has entered a stage of practical application in a wide range of biomedical research fields.

Looking back at the breakthrough in identification depth of single-cell transcriptome sequencing technology ten years ago^45-48^, the Smart-seq-based single-cell RNA-seq technique could identify around 30,000 transcripts from one human tumor cell^45^, accounting for 13% of the total human transcripts (ca. 230,000^49^). In this study, up to 3,000 proteins quantified in one human tumor cell account for 15% of the total human proteins (ca. 20,000^49^), reaching the similar level of single-cell RNA-seq technique at that time. Considering the explosive development of the single-cell transcriptome sequencing technology over the past decade^50-52^, we may now be on the eve of or even in the outbreak of the single-cell protein sequencing technology based on the shotgun proteomics strategy. In the near future, the identification depth and throughput of single-cell proteomic analysis will be further improved, reaching the level of practical and popular application. In addition, by adopting the microfluidic sample pretreatment technique, it will be possible to combine it with single-cell genome, transcriptome, and metabolome analysis technologies to form a “true” single-cell multi-omics analysis technology for single-cell individuals.

These will undoubtedly bring unprecedented powerful tools for people to understand the variations of cellular heterogeneity in life activities.

## Methods

### Cell culture

HeLa, A549 and U2OS cells were used in the experiments. The HeLa and U2O2 cells were maintained in DMEM with high glucose/pyruvate (Invitrogen) supplemented with 10% FBS (Gibco). The A549 cells were maintained in F-12K supplemented with 10% FBS. All cell lines were maintained in a 5% CO_2_ incubator at 37 °C. For single-cell analysis, cells in a 6 cm dish were collected at 60-80% confluency using 0.25% trypsin with 0.02% EDTA, and washed five times with PBS.

### Single cell capture

We used the liquid handling robot modified from a system for single-cell soring^26^ to achieve the operations of single cell capture and sample pretreatment. It consisted of a microscopic imaging module, a capillary probe (10 cm length, 250 μm o.d., 150 μm i.d., tip size, 35 μm i.d.) connected with a high-precision syringe pump (1701N, Hamilton), an automated *x-y-z* translation stage, and a system control module. For cell capture, 2 mL of cell PBS suspension with a cell density of ca. 1×10^3^ cells/mL was added in a petri dish (35 mm diameter) fixed on the translational stage of the robot. The microscopic imaging module first took bright-field or fluorescence image of the cells in the cell suspension settled on the bottom of the petri dish. The target cells in the image were selected, and their location coordinates were automatically calculated by the control module. Then the tapered tip of the capillary probe was controlled to automatically align the target cell by moving the translational stage and suck it into the probe by aspirating 15 nL of cell suspension into the probe by the syringe pump. After the target cell was picked up by the capillary probe, it was then deposited to the bottom of an insert tube.

### Single-cell sample pretreatment

After picking up the single target cell into the insert tube, we used the robot to perform the subsequent sample pretreatment operations, including cell lysis, protein reduction, alkylation, enzymatic digestion and termination of the digestion reaction, by sequentially adding 100 nL of RapiGest SF (Waters), tris (2-carboxyethyl) phosphine (TCEP), iodoacetamide (IAA), mixed enzyme (Lys-C and trypsin), and formic acid (FA) solutions to the insert tube, respectively. To avoid cross-contamination between different reagents and samples, the capillary probe was washed with water three times before aspirating new reagents or samples. The sample vials with the insert tubes were placed in the sample tray of the LC autosampler for LC-MS/MS analysis.

### LC-MS/MS analysis

A capillary LC column was used in the LC separation of the digested peptide samples. The single-cell samples in the insert tubes were injected by an autosampler coupled to an EASY-nLC 1200 LC (ThermoFisher Scientific).

The separated peptide components of single cell samples were detected by a trapped ion mobility-time of flight mass spectrometer (timsTOF Pro, Bruker). Data dependent acquisition (DDA) mode and data independent acquisition (DIA) mode were both used in the research. For DDA mode, the parallel accumulation serial fragmentation (PASEF) acquisition mode was used. The precursor ions obtained by the primary mass spectrum underwent secondary fragmentation through collision induced dissociation (CID). The conditions of the DIA mode was set on the basis of the DDA results.

### Raw data analysis

The DDA raw files were analyzed with Spectromine software (version3.2, Biognosys AG, Schlieren, Switzerland) using default settings. Protein label-free quantitative identification from the DDA dataset was searched against the SwissProt/UniProt human proteome database (UP000005640_9606.fasta, Homo Sapiens: 20,614 entries). Enzyme specificity was set to trypsin cleaving C-terminal to arginine and lysine. A maximum of one missed cleavage were allowed. Cysteine carbamidomethylation was set as a static modification, acetylation on protein N-terminus and oxidation on methionine set as variable modifications. The peptide-spectrum matches (PSM), peptides and protein groups were filtered at 1% FDR. The DIA raw files were analyzed with DIA-NN software (version 1.8) in library-free search mode using default settings. False discovery rates (FDR) were controlled at 1% on precursor level.

### Data analysis and visualization

Most data were analyzed by R (version R4.13) and visualized by R package ggplot2 (version 3.3.5). The uniform manifold approximation and projection (UMAP) analysis was carried out by the umap-learn module in Python3. Differential analysis was carried out using t test (unpaired) in the stats package. Hierarchical cluster analysis (HCA) was visualized by R package pheatmap (version 1.0.12). Gene Ontology (GO) enrichment was carried out by the goatools module (v1.0.15) in Python3. Kyoto Encyclopedia of Genes and Genomes (KEGG) enrichment was carried out by the kobas module (version 3.0) in Python3. The interaction network map of differential proteins was analyzed on String database (version 11.5).

## Acknowledgement

This article is to commemorate Professor Peng-Yuan Yang, one of the pioneers of proteomic research in China, and we are grateful for his forward-looking suggestion to initiate our first attempt in single-cell proteomic analysis more than 10 years ago. We thank Professor Catherine C. L. Wong (Peking Union Medical College Hospital) and Dr. Zi-Yi Li (Zhejiang University) for their help with training in the use of LC-MS/MS systems in the early stage of our work. We thank Dr. Cheng-Pin Shen (Shanghai Omicsolution Co.) for his help in data analysis. The authors acknowledge the financial support provided by National Natural Science Foundation of China (Grants 21827806 and 21435004), and National Ministry of Science and Technology (Grant 2021YFA1301601).

## Author contributions

Q.F., Y.W. and Z.Y.G. conceived the idea. Y.W. and H.F.W. built the instrument. Y.W. and H.F.W. wrote the software. Z.Y.G. and Q.W. cultured the cells. S.W.S., Z.Y.G., Y.R.J. and Q.W. explored and optimized the LC-MS conditions. Y.W. and Z.Y.G. planned and performed experiments. Y.R.J., Q.W., S.W.S. and W.X.Y. provided nano-LC columns. Y.W., Z.Y.G., L.Z., Y.R.J., J.W., Q.W., J.B.C., W.X.Y., Q.Q.X., Q.X.F. and Q.F. contributed to data analysis and interpretation, as well as figure preparation. Q.F., Y.W., Z.Y.G. and J.B.C. wrote the manuscript. Q.F. and J.Z.P. supervised the studies and acquired funding.

