## Supplemental Figure 1 and 2 for "Pick-up Single-Cell Proteomic Analysis for Quantifying up to 3000 Proteins in a Tumor Cell"

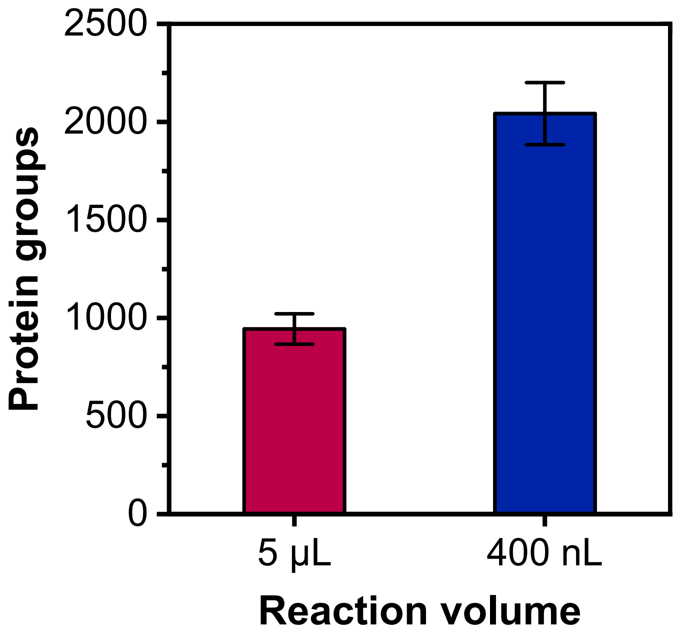


**Supplementary Figure 1.** Comparison of the protein group number quantified in single HeLa cells with microliter- and nanoliter-scale reaction volumes for sample pretreatment. In average, 944 ± 77 ($n$ = 4) and 2042 ± 158 ($n$ = 4) protein groups were quantified under the DDA mode with the reaction volumes of 5 μL and 400 nL, respectively.

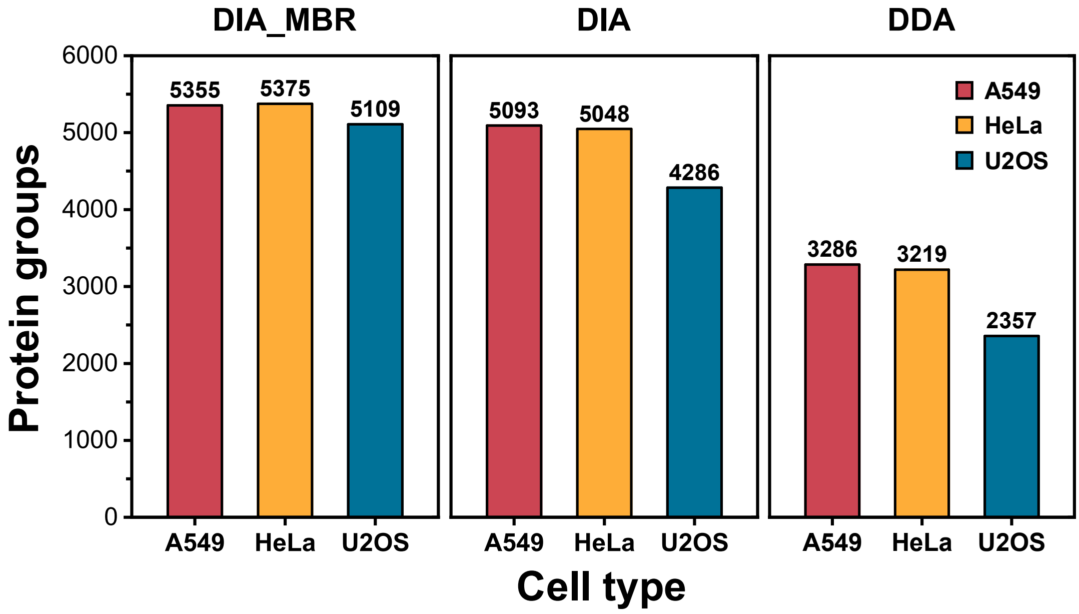


**Supplementary Figure 2.** Cumulative total number of protein groups quantified in single tumor cells. Under the DIA mode with MBR, total numbers of 5355, 5375 and 5109 protein groups were quantified cumulatively from single A549 ($n$ = 37), HeLa ($n$ = 44) and U2OS ($n$ = 27) cells, respectively. Under the DIA mode without MBR, total numbers of 5093, 5048 and 4286 protein groups were quantified for the same A549, HeLa and U2OS cell samples, respectively. Under the DDA mode, total numbers of 3286, 3219 and 2357 protein groups were quantified from A549 ($n$ = 56), HeLa ($n$ = 68) and U2OS ($n$ = 24) cells, respectively.
